# Depression symptoms are associated with affective neural processing during sleep and rest

**DOI:** 10.64898/2026.03.03.709353

**Authors:** Xuanyi Lin, Nicholas J Lew, Matthew Cho, Ken A. Paller, Eitan Schechtman

## Abstract

Sleep supports offline information processing and is essential for cognitive and emotional functioning. Abnormal sleep patterns are a hallmark of affective disorders. We hypothesized that affective symptoms occur with maladaptive neural processing during offline periods. To test this idea, we used multivariate EEG decoding with cross-state classification. A model trained on EEG data acquired while participants (*N* = 52) viewed emotional images was used to classify stimulus valence. Applying this model to data collected during a nap revealed the re-emergence of affective neural patterns. Critically, offline reinstatement of patterns reflecting negatively valenced processing predicted greater depressive symptoms across participants. These associations reflected both cognitive-affective and somatic-performance depression subscales, and they generalized across sleep stages and wakeful rest. The finding that offline neural information processing is linked with emotional well-being supports a model whereby maladaptive negative biases can be perpetuated during rest, potentially shaping the progression of affective disorders.

## Introduction

Sleep and wakeful rest play a critical role in the processing of newly acquired memories^1,2^. Neural traces supporting recently encoded information are spontaneously re-expressed during sleep, leading to memory consolidation and reorganization^3,4^. Whereas the mechanisms supporting sleep’s active contribution to memory have been systematically explored, far less is known about the mechanisms supporting sleep’s role in shaping affective states. Growing evidence suggests that affective neural networks remain engaged and activated during sleep, shaping waking emotional function. Emotional memories, for example, are preferentially consolidated compared to neutral ones, particularly when emotional salience is high^5,6^.

Importantly, the intersection of sleep and affective states is not limited to its contribution to memory. At the systems level, sleep produces measurable changes in the functional organization of affective brain circuits. Neuroimaging studies show that sleep loss and recovery bidirectionally modulate amygdala responsivity to emotional stimuli and its functional connectivity with medial and dorsolateral prefrontal cortex, a circuit central to emotional evaluation and regulation^7–9^. These local circuit effects are embedded within broader sleep-dependent changes in large-scale brain network dynamics supporting affective processing^10^. These modulations bear measurable behavioral benefits: sleep has been shown to improve mood^11,12^, reduce anxiety symptoms^13^, and enhance emotional control^9^. The contribution of sleep to emotional states, both in the short and long term, have been shown to relate to dream contents^14^. These findings align with theoretical models proposing a central role for sleep in shaping affective states and guiding emotional regulation^15^.

Studies directly examining mechanisms of affective processing during sleep are scarce. Recent work has shown that the sleeping brain continues to differentiate emotional valence, exhibiting distinct neural responses to positive and negative stimuli even in the absence of conscious awareness^16^. Other studies have examined spontaneous and triggered patterns of neural reactivation during sleep as they relate to affective processing^17–19^. These observations further demonstrate that sleep involves processing by neural affective networks, yet leave open the question of the relationship between these activation patterns and waking well-being. Critically, it remains unclear whether affective processing during sleep is linked to symptoms experienced by people with affective disorders such as major depressive disorder (MDD).

Two frequent symptoms of MDD are insomnia (difficulty falling asleep) and, alternatively, hypersomnia (difficulty remaining awake)^20^. Furthermore, sleep continuity is compromised in MDD^21^ and the duration of rapid-eye-movement sleep (REM) is increased^21,22^, mirrored by a decrease in slow-wave sleep (SWS), the deepest stage of sleep^21,23^. Although there is some evidence for abnormal neural activation during sleep in people with MDD (e.g., increased activation in the amygdala and anterior cingulate cortex during REM sleep^24^), neural activation patterns during sleep have not been thoroughly studied. Cognitive models of depression emphasize persistent negative biases in attention, appraisal, and memory, which contribute to symptom maintenance^25,26^. While these biases in cognitive processing are well documented during wakeful cognition, it is unknown whether they persist into sleep, shaping the spontaneous neural landscape in offline brain states. If negative biases do perpetuate during offline processing, they may contribute to the progression of MDD. Furthermore, affective traits (e.g., depression symptoms) and states (e.g., moods) may differ in how they are impacted by processing during sleep. Depressive symptoms are relatively stable over time and may reflect enduring biases in emotional representation, whereas momentary mood states fluctuate over shorter timescales and may be more sensitive to contextual and physiological factors associated with sleep. Distinguishing between trait-like and state-dependent influences on affective processing during sleep is therefore essential for understanding the functional role of sleep in emotional health.

In the present study, we tested whether valence-driven neural patterns decoded from wake EEG re-emerge during a daytime nap, and whether these activation patterns relate to individual differences in affective traits, mood states, and post-sleep performance on an emotional memory task (Fig 1). High-density EEG was recorded as participants viewed affective stimuli (positive vs negative images, Fig 1c), enabling time-resolved multivariate decoding of emotional valence from scalp-recordings. These wake-trained decoding models were then applied to continuous EEG recorded during a 90-min nap to identify spontaneous activity patterns linked with positively valenced and negatively valenced processing. We examined how these valence-based patterns related to (i) trait depressive symptomatology, (ii) pre-nap mood state and post-nap mood change, and (iii) post-sleep emotional memory. In addition, valence-related activation patterns expressed during sleep were compared with those observed during pre-sleep quiet rest, allowing us to distinguish neural processes that are sleep-specific from those also occurring during wakeful rest. By integrating multivariate neural decoding, high-density EEG, behavioral measures, and an individual-difference approach, we demonstrate that neural affective activity patterns during rest are linked to waking well-being.

**Figure 1.**
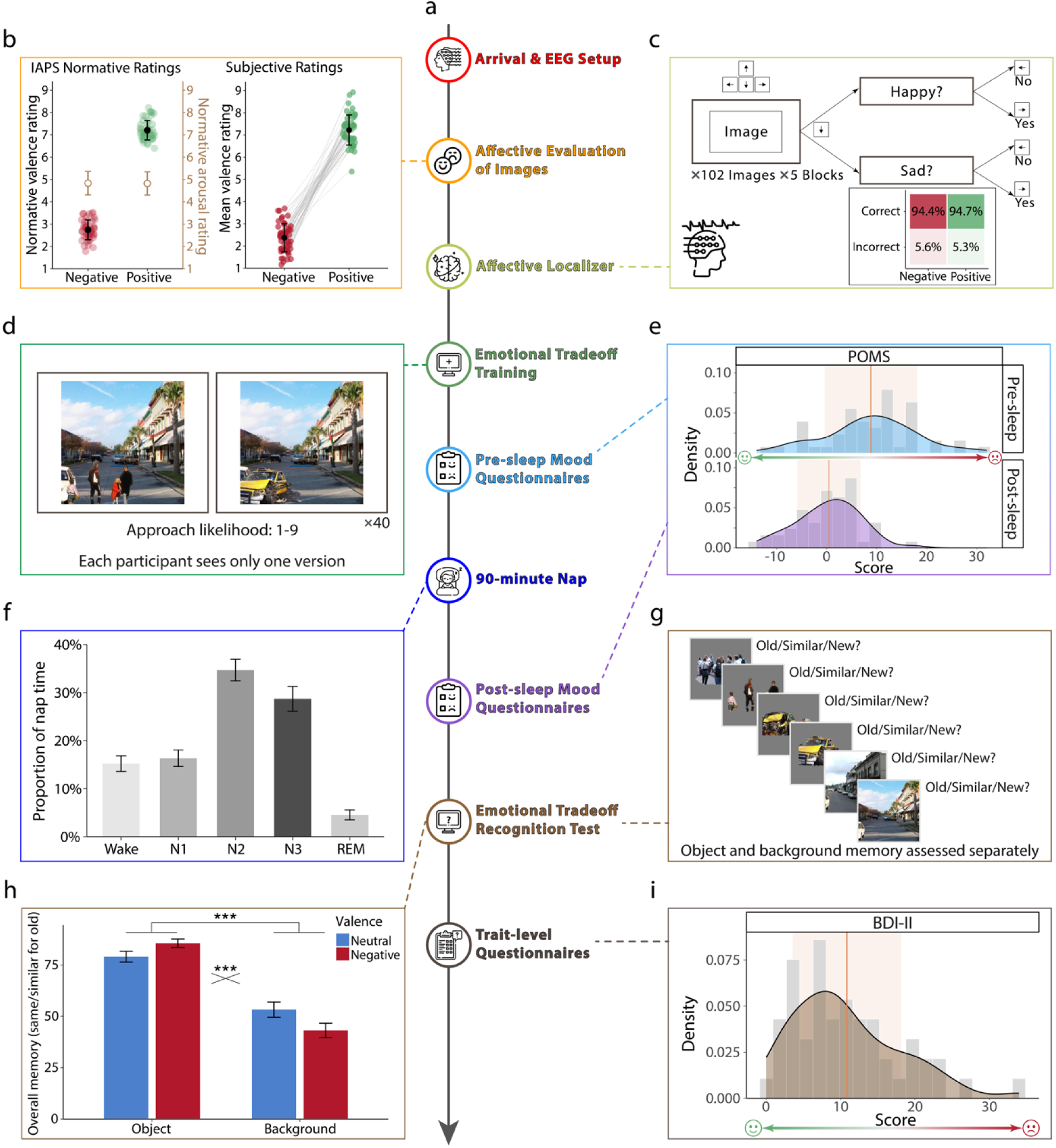
Experimental design and behavioral results. (a) Vertical experimental timeline illustrating the study procedure. In a single afternoon session, participants completed an affective image evaluation task and an affective localizer task while high-density EEG data was recorded. Then, they trained on an emotional memory task and completed a mood assessment before sleeping in the lab for 90 min. After waking up, they completed a mood assessment, a memory test, and trait-level affective questionnaires. Dashed lines connect each step to the corresponding panel. (b) Affective evaluation of images from the International Affective Picture System (IAPS). Left: IAPS normative rating show that negative and positive images differed in valence (red/green circles), but not in arousal ratings (brown circles) at the image level. Error bars reflect SD. Right: subjective valence ratings, averaged within participant for negative and positive stimuli (red/green circles, respectively), showing reliable differentiation between the two classes. Lines connect within-subject ratings across valence categories. Error bars reflect SD. (c) Schematic of the affective localizer task. Participants viewed 102 images in each of five blocks and reported each image’s valence by responding to an on-screen prompt (“Happy?”/ “Sad?”) using the keyboard. Inset: confusion matrix summarizing classification performance of behavioral responses (% correct and incorrect). (d) Example stimuli from the training phase of the emotional tradeoff memory task. Scenes consisted of neutral backgrounds paired with either neutral or negative objects (left and right, respectively). Participants rated their likelihood to approach the scene. Each participant viewed only one version of each scene. (e) Distribution of pre-sleep and post-sleep mood state, measured by the Profile of Mood State questionnaire (POMS). Density plots depict score distributions; vertical red lines reflect mean; shaded regions indicate ±1 SD. Mood improved following sleep. (f) Proportion of nap time spent in each sleep stage during the 90-min nap opportunity. Bars indicate mean ± SD. REM – rapid-eye-movement sleep; N1/N2/N3 – stages 1/2/3, respectively. (g) Schematic of the recognition test for the emotional tradeoff memory task. Object and background memory were assessed separately using old/similar/new judgments. (h) Results for the emotional tradeoff memory task. Recognition performance for objects and backgrounds as a function of valence. Negative scenes enhanced object memory but reduced background memory relative to neutral scenes (valence × object/background interaction). Bars indicate mean ± SD. *** - p < .001. (i) Distribution of trait-level depressive symptom severity (Beck Depression Inventory-II; BDI-II). Density plot illustrates score distribution; vertical red line reflects mean; shaded region indicates ±1 SD.

## Results

Fifty-two healthy young participants were recruited from the campus community. In the lab, they engaged in multiple tasks and questionnaires and were given a 90-minute nap opportunity, all while high-density EEG data were collected (Fig 1a). They were first asked to rate the valence for a set of negatively and positively valenced images. Their ratings confirmed that they viewed the negatively valenced subset as more negative than the positively valenced subset (*t*(51) = 30.89, *p* < .001; Fig 1b, right). Next, participants completed an affective localizer task, which was used to train the EEG classifier. Participants viewed the same images and indicated their valence (i.e., positive or negative; Fig 1c). Performance was close to ceiling for both stimulus categories (negative: 94.4% correct; positive: 94.7% correct). A complementary signal-detection analysis confirmed high sensitivity (median d’ = 3.66, mean ± SD = 3.65 ± 0.96). Before napping in the lab, participants filled out questionnaires examining their mood (Profile of Mood State; POMS^27^), anxiety levels (State-Trait Anxiety Inventory, State questions only; STAI-S^28^), and sleepiness (Stanford Sleepiness Scale; SSS^29^). These questionnaires were completed again after the nap. On average, participants slept for 85.8% of the total time in bed and spent time in all sleep stages (see Fig 1f, Supp Fig 1, and Supp Table 1).

Compared to pre-nap POMS (mean ± SD = 8.83 ± 9.24; higher values indicating more negative mood), post-nap POMS (mean ± SD = 0.42 ± 6.35) significantly decreased (*t*(51) = 7.10, *p* < .001, Fig 1e), suggesting improvement in mood over the nap period. Similarly, post-nap STAI-S (mean ± SD = 30.60 ± 5.79; higher values indicating more anxiety) was significantly lower than pre-nap STAI-S (mean ± SD = 33.87 ± 7.53, *t*(51) = 3.70, *p* < .001), indicating a decrease in anxiety level. Subjective sleepiness also significantly decreased following sleep (pre-nap SSS mean SD = 4.10 1.30, post-nap SSS mean SD = 2.60 0.774, ΔSSS mean SD = -1.50 1.23, *t*(51) = -8.80, *p* < .001). Collectively, these findings are consistent with prior reports demonstrating improvements in mental well-being following daytime sleep^30,31^. After sleep, participants filled out trait level questionnaires. On average, participants reported low to moderate depressive symptoms (Beck Depression Inventory-II; BDI-II^32^, mean SD = 10.83 7.30, range = 0–34; Patient Health Questionnaire; PHQ^33^, mean SD = 5.73 4.55, range = 0–18; higher values reflect greater severity of depressive symptoms), consistent with a non-clinical sample (see Supp Table 2 for a summary of all the questionnaires).

To establish a reliable method for monitoring affective neural states, we first examined whether emotional information could be reliably decoded from neural responses collected during the affective localizer task (Fig 2a). Using multivariate pattern classification, a linear support vector machine was trained at each time point on EEG scalp patterns to discriminate positive vs. negative images in a cross-validated, time-resolved manner (Fig 2a). This time-resolved decoding revealed that the emotional valence of the presented stimuli could be discriminated from EEG scalp patterns significantly above chance (50%) over two time windows: -108–1498 ms (*p_corrected_* < .001) and 1564–1784 ms (*p_corrected_* = .006, Fig 2b), indicating that valence-related information is decodable from EEG data across an extended post-stimulus period. Note that decoding estimates were computed using 102-ms sliding windows (22-ms step size); thus, each time point reflects information integrated across a ±51-ms interval around the window center. The cluster onset in the late pre-stimulus period should therefore be interpreted cautiously and likely reflects temporal smoothing or slowly varying baseline rather than stimulus-evoked processing per se. Models trained at each time point did not generalize to other time points, suggesting that different neural patterns drove the classifiers’ performance throughout the time course of the trial (Supp Fig 2). Accuracy peaked at 398 ms (mean ± s.e.m. = 66.35 ± 0.017%). Classifier accuracy (measured as areas under the curve during significant periods, i.e., -108–1498 and 1564–1784 ms) was not correlated with any of the affective trait measures across participants (e.g., BDI-II, *p*s > .153; correlation for BDI-II shown in Figure 2c).

**Figure 2.**
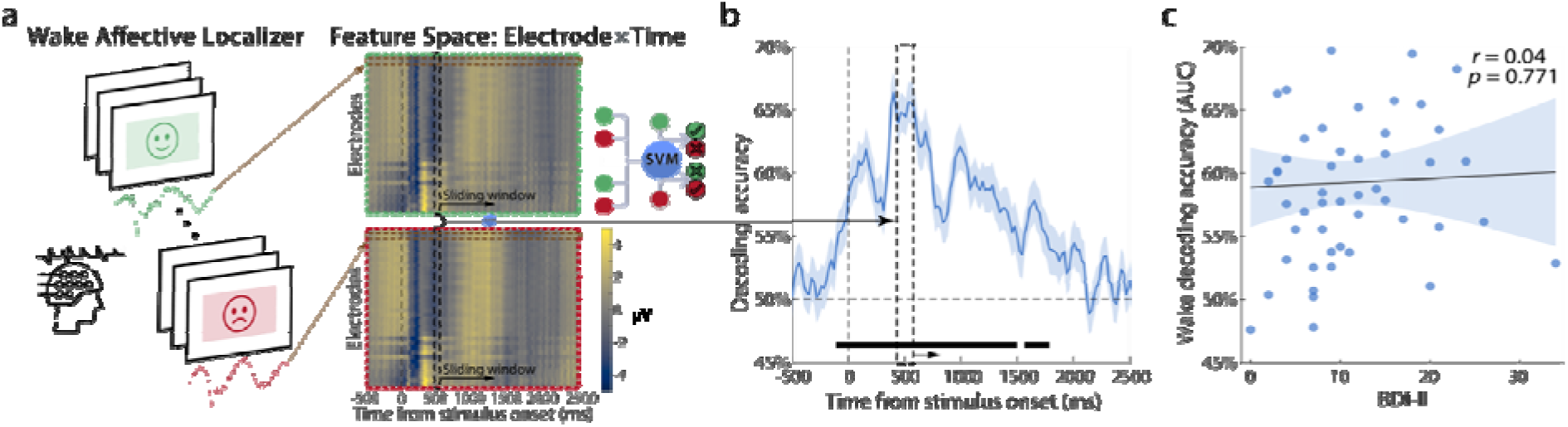
EEG neural patterns reflect the valence of affective stimuli during wakefulness. (a) Schematic of the wake affective localizer and decoding approach. At each time point relative to image onset, multichannel EEG scalp voltages were used as features to train a linear support vector machine (SVM) classifier across trials. The classifiers classified image valence in a time-resolved, cross-validated manner (inset). Green- and red-framed boxes reflect responses following presentation of positive and negative images, respectively. Data shown for a single representative participant. (b) Group-averaged decoding accuracy time course (mean s.e.m.), aligned to stimulus onset. Emotional valence was decodable above chance (50%; dashed line) across a broad temporal window, with significant clusters indicated by the horizontal bar (two clusters: -108–1498 ms, p_corrected_ < .001; 1564 to 1784 ms, p_corrected_ = .006). (c) Wake decoding accuracy (area under the curve throughout both significant periods; AUC) was not significantly correlated with depressive symptom severity, as measured by Beck Depression Inventory-II (BDI-II) scores.

To test whether affective neural states re-emerge during sleep, we applied the wake-trained classifiers to the EEG data collected during the nap in a cross-state decoding framework (Fig 3a). Specifically, we applied the classifiers trained at each time point relative to stimulus onset during wakefulness to the continuous EEG data recorded during the nap. This yielded an assigned label (positive or negative) for each time-specific classifier trained during wakefulness and each data segment collected during sleep. For each of these wake-trained classifiers, we aggregated the assigned labels across the entire nap, independent of sleep stage, by averaging the classifier output over the full duration of the nap (Fig 3a, right panel; see Supp Fig 3a for breakdown by sleep stage). This analysis yielded, for each participant, a profile of affective activation pattern collapsed over the entire nap, indexed by the time point at which classifiers were trained during wake (Fig 3b, third panel). This wake-training-time-resolved profile therefore indicates which wake-evoked affective representations – defined by their latency during wake – were preferentially re-expressed during the nap (note that classifiers trained on different time points reflected distinct valence-sensitive neural patterns; Supp Fig 2).

**Figure 3.**
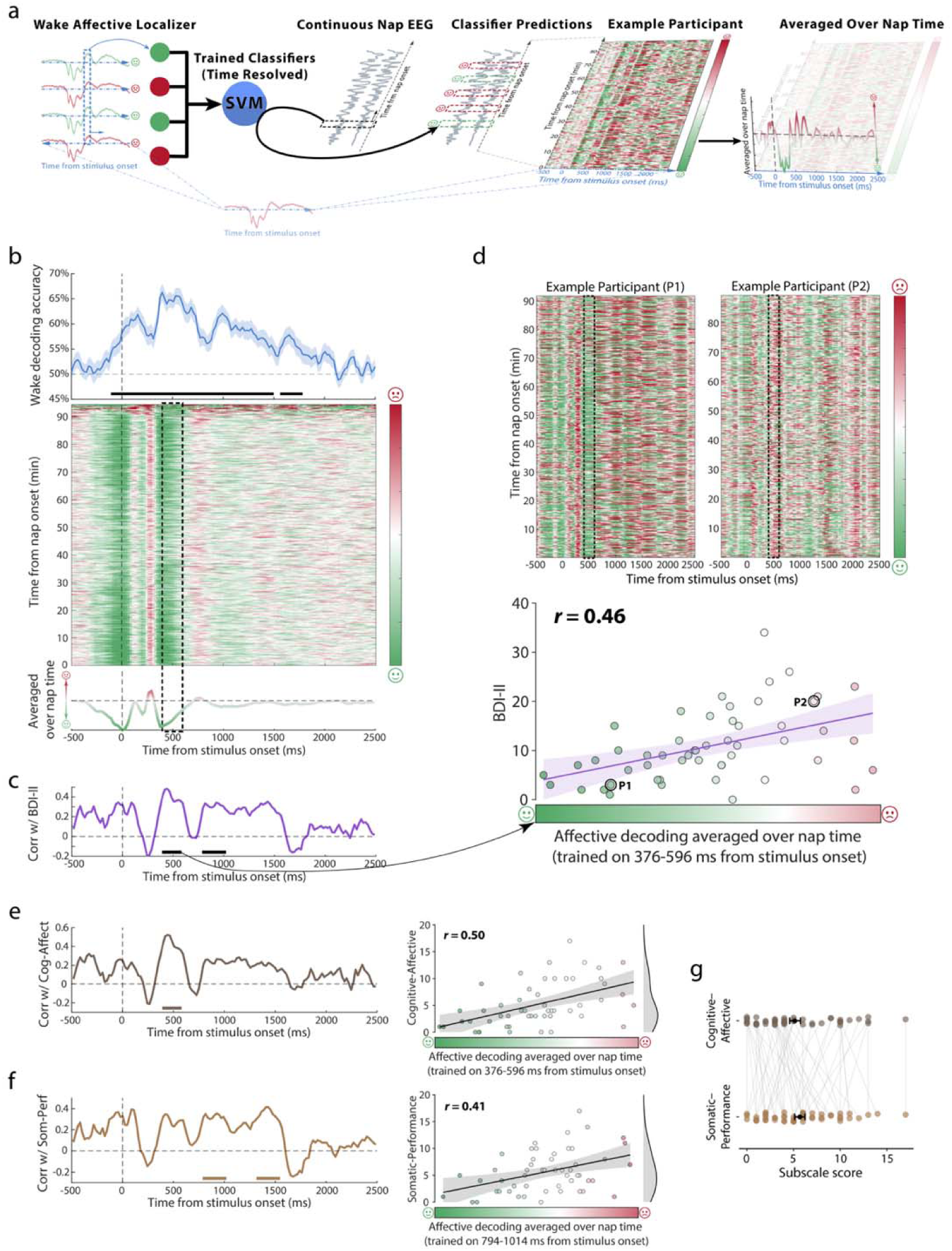
The re-emergence of affective representations during sleep is associated with depressive symptoms. (a) Schematic of the cross-state decoding framework. Time-resolved classifiers were trained on EEG data from the wake affective localizer at each time point relative to stimulus onset, such that each classifier indexed the neural representation of affective processing at a specific post-stimulus latency during wake (see Fig 2). These wake-trained classifiers were then applied to continuous nap EEG, yielding time-resolved predictions of affective activation throughout sleep. Classifier outputs were averaged across the full nap duration to produce, for each participant, a wake-trained-time-resolved profile of affective reactivation during sleep. Red/green – negative/positive valence, respectively. (b) Wake decoding accuracy (top, same as Fig 2b) and corresponding cross-state decoding results (middle) averaged across participants. The heatmap depicts classifier predictions across nap time (y-axis) as a function of wake training time (x-axis), illustrating that affective representations defined during wake are expressed throughout sleep. Averaging predictions across nap time yields a wake-time-resolved affective activation profile (bottom). The dashed box in the middle panels reflects the time span of the early cluster (see panel c), between 376–596 ms. (c) Time-resolved correlations between affective reactivation during sleep and depressive symptoms (measured using BDI-II) are shown below, with significant clusters (corrected for multiple comparisons) indicated by horizontal lines. (d) Example cross-state decoding heatmaps from two representative participants, illustrating individual variability in the expression of wake-defined affective representations during sleep. The dashed box reflects the first cluster, as in panel b. The scatter plot shows the relationship between BDI-II symptoms and mean affective activation during sleep, averaged across the early significant wake-time window (376–596 ms). Data for the representative participants is marked as P1 and P2. (e-f) Left – Time-resolved correlations between affective activation during sleep and cognitive–affective (e) and somatic–performance (f) depression subscales, plotted as a function of wake training time. Right – corresponding scatter plots for significant wake-time windows. (g) Distribution of cognitive–affective and somatic–performance subscale scores across participants, illustrating variability in affective symptomatology. Black dots and error bars mark mean ± s.e.m.

The assigned labels essentially reflect similarities between affective neural patterns established during wakefulness and neural activity during sleep. Critically, however, the functional relevance of these cross-state pattern reinstatements remains unclear, as there is no ground truth to which the assigned values can be linked – in many decoding applications, classifier outputs can be validated against known labels (e.g., stimulus category or behavioral responses), allowing direct assessment of predictive accuracy. In contrast, sleep provides no such externally verifiable labels, and therefore the functional significance of cross-state pattern reinstatement cannot be established through accuracy metrics alone. The final inferential step of our approach was therefore the most critical one: we examined whether re-emergence of affective neural patterns during sleep was correlated with trait- and state-level affective metrics. This was done by correlating the wake-training-time-course of affective activation during sleep with behavioral measures using a time-resolved, cluster-based permutation approach, which corrects for multiple comparisons and avoids spurious correlations (Fig 3b, bottom).

We first examined depressive symptom severity, indexed by BDI-II scores, and found that it was positively associated with the re-emergence of neural patterns linked with processing negatively valenced information. Two discrete clusters in post-stimulus intervals during wakefulness were significantly correlated with BDI-II (Fig 3c): one observed for classifiers trained on 376–596 ms relative to stimulus onset (*p_corrected_* = .019; *r* = 0.46, *p_uncorrected_* < .001, when affective decoding is averaged over the cluster time window, Fig 3d), and the other observed for classifiers trained on 794 –1014 ms relative to stimulus onset (*p_corrected_* = .049; *r* = 0.39, *p_uncorrected_* = .004). Note that all clusters fall within the timespan of above-chance classification during wakefulness (−108–1498 ms and 1564–1784 ms), with the first cluster coinciding with the peak accuracy (∼398 ms). These correlations were not driven by differences in wake classification accuracy, as the correlations between BDI-II and the accuracy for these two time windows during wake were not significant (*r*s < 0.25, *p_uncorrected_*s > 0.0759). These results indicate that individuals with more severe depressive symptoms exhibited a stronger bias toward negative-valence neural activation patterns during the 90-minute nap.

BDI-II comprises two well-characterized subscales that capture partially dissociable dimensions of depressive symptomatology: a cognitive-affective (Cog-Affect) subscale reflecting negative mood, self-evaluation, and affective distress (e.g., sadness, pessimism, guilt), and a somatic-performance (Som-Perf) subscale reflecting bodily and motivational symptoms such as fatigue, sleep disturbance, and reduced activity levels^32,34^. To further explore the relationship between depressive symptoms and affective activation during sleep, we separately examined the two subscales of BDI-II (Cog-Affect, mean ± SD = 5.15 ± 4.18; Som-Perf, mean ± SD = 5.67 ± 3.92, Fig 3g). We used the same time-resolved, cluster-corrected correlation approach, this time correlating the time-series data with each subscale separately. Distinct temporal clusters emerged from this analysis, mapping on to the two clusters identified for the full BDI-II scale. When examining the cognitive-affective subscale, an early cluster (376–596 ms) emerged (*p_corrected_* = .008; *r* = 0.50, *p_uncorrected_*< .001, Fig 3e), whereas the later cluster emerged selectively for the somatic-performance subscale (794–1014 ms, *p_corrected_* = .039, *r* = 0.41, *p_uncorrected_* = .003, Fig 3f). A third cluster, which was not observed when considering the full BDI-II scale, was also selectively correlated with the somatic-performance subscale (1300–1542 ms, *p_corrected_* = .016, *r* = 0.43, *p_uncorrected_*= .001). Together, these findings suggest that temporally distinct wake-evoked affective representations re-emerged during sleep, and their expression differentially tracked cognitive-affective vs. somatic-performance components of depressive symptomatology, indicating a dissociation in how specific facets of depression relate to affective processing during sleep.

We next assessed whether the depressive symptom effect differed across sleep stages. When cross-state decoding was computed separately for each sleep stage, significant correlations were observed for a similar early wake-training interval (∼376–596 ms) during Stages 1 (*p_corrected_* = .019, during 376–596 ms; *r* = 0.47, *p_uncorrected_* < .001), 2 (*p_corrected_* = .030, during 376–596 ms; *r* = 0.45, *p_uncorrected_* < .001), and 3 (*p_corrected_* = .040, during 420–596 ms; *r* = 0.44, *p_uncorrected_* = .002) of sleep, as well as during pre-sleep or mid-sleep wakefulness periods (*p_corrected_* = .031, during 376–574 ms; *r* = 0.41, *p_uncorrected_* = .003; Supp Fig 3b). During rapid-eye-movement (REM) sleep, no significant clusters overlapped with the early time cluster, but this may be due to the fact that only 21 of the 52 participants entered REM during their nap, as is typical for afternoon naps, limiting statistical power. This stage-general expression indicates that trait-like depressive symptoms are associated with a pervasive bias toward negative-valence activation patterns across the sleep cycle, rather than being confined to a specific sleep stage. None of the sleep stages had significant temporal clusters that overlapped with the later cluster (∼794–1014 ms; *p*s*_corrected_* > .062). A late cluster (1586–1762 ms), significant only during REM, was found to be negatively correlated with BDI-II scores (i.e., more negative scores predicted fewer symptoms; *p_corrected_* = .048; *r* = -0.62, *p_uncorrected_* = .003, Supp Fig 3b). However, as previously mentioned, this analysis utilized data from a smaller sample size (*n* = 21 who entered REM sleep).

We next examined whether the two BDI subscales were associated with sleep-stage-specific affective activation. For the cognitive-affective subscale, significant clusters that overlap with the early wake-training interval (∼376–596 ms) were observed during Stages 1 (*p_corrected_* = .009, during 376–596 ms; *r* = 0.51, *p_uncorrected_* < .001), 2 (*p_corrected_* = .008, during 376–596 ms; *r* = 0.50, *p_uncorrected_* < .001), and 3 (*p_corrected_* = .020, during 398–574 ms; *r* = 0.47, *p_uncorrected_* < .001) of sleep, as well as during pre-sleep or mid-sleep wakefulness periods (*p_corrected_* = .026, during 376–574 ms; *r* = 0.43, *p_uncorrected_* = .002; Supp Fig 3c). No significant correlation was observed during REM sleep in this time window. For the somatic-performance subscale, the analysis yielded significant clusters which overlapped with the later cluster (∼794–1014 ms) during Stage 2 of sleep (*p_corrected_* = .012, during 772–1036 ms; *r* = 0.46, *p_uncorrected_* < .001) and pre-sleep or mid-sleep wakefulness (*p_corrected_* = .029, during 772–1014 ms; *r* = 0.43, *p_uncorrected_* = .001; Supp Fig 3d). A late cluster (1608–1762 ms) showed a negative correlation between REM sleep and somatic-performance subscale (*p_corrected_* = .041; *r* = -0.65, *p_uncorrected_*= .002; Supp Fig 3d), suggesting that the negative correlation with REM observed for the full BDI-II scale was driven by the somatic-performance subscale.

We next turned to additional behavioral measures using the same analytic approach, considering both the full nap and individual sleep stages. Across the nap, pre-sleep mood state scores, as measured using POMS, were correlated with negative activation for classifiers trained on an overlapping early time window 376–618 ms (*p_corrected_* = .026, *r* = 0.40, *p_uncorrected_*= .003; Fig 4a and Supp Fig 4a). This indicates that participants entering sleep in a more negative mood showed greater negative emotional bias in their spontaneous affective activation patterns during nap. This result was not driven by differences in wake classification accuracy (*r* = -0.17, *p* = .229), but may be at least partially driven by the strong correlation between BDI-II and pre-sleep POMS (*r* = 0.58, *p* < .001; see Supp Table 3 for correlations between BDI-II and other behavioral measures). Nevertheless, sleep-stage-specific analyses revealed a different pattern from that observed for depressive symptoms. We only found a significant cluster spanning the range between 420–618 ms during Stage 3 of sleep (*p_corrected_* = .026; *r* = 0.41, *p_uncorrected_* = .003; Supp Fig 3e). We also examined whether neural affective patterns observed during sleep were associated with post-sleep mood or mood change, as indexed by post-nap POMS scores and post-minus-pre nap POMS differences. Using the same time-resolved, cluster-based correlation approach, no robust clusters survived correction for multiple comparisons (*p_corrected_*s > .125), indicating that sleep-based affective activation was not strongly associated with immediate post-sleep mood outcomes (Supp Fig 4b-c).

**Figure 4.**
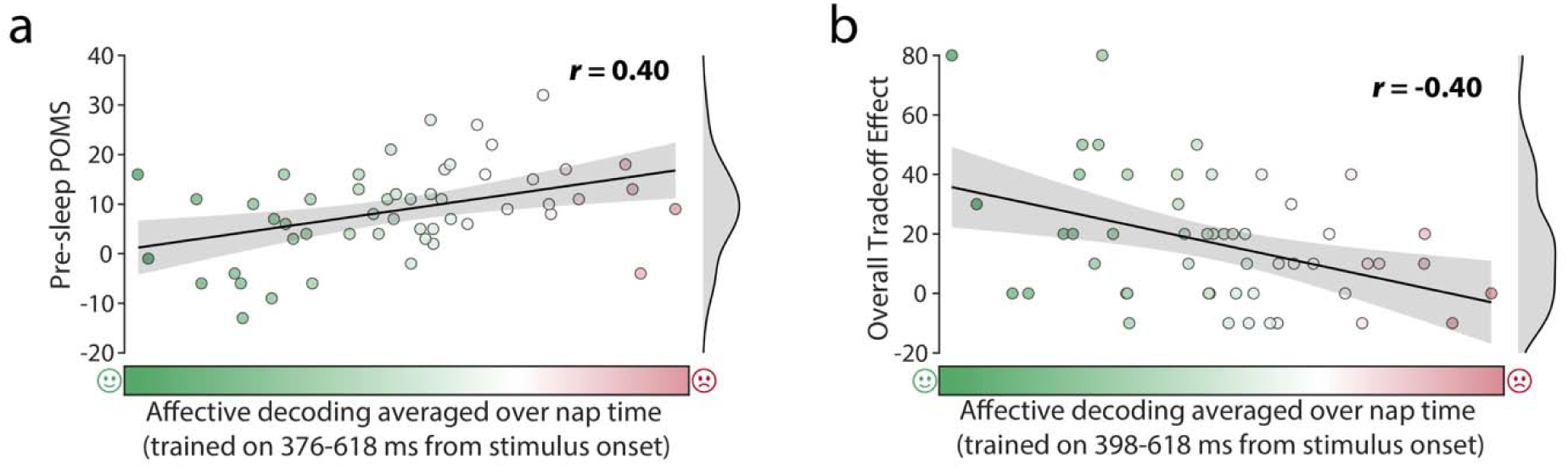
Affective activation during sleep was positively associated with pre-sleep mood (a) and negatively associated with memory performance in the emotional tradeoff memory task (b). POMS – Profile of Mood State questionnaire.

Previous studies have shown an emotional tradeoff memory effect, whereby negative objects are remembered better than neutral ones, whereas the contextual background of the negative objects are remembered worse than the contexts of their neutral counterparts^35,36^. This effect has been shown to be enhanced by sleep relative to wakefulness^5,6^. We incorporated this memory task in our protocol to examine whether tradeoff effects are linked with affective activation patterns during sleep. We first examined whether an overall trade-off effect emerged in our data. Overall recognition accuracy in the post-nap emotional tradeoff test (Fig 1h) was analyzed using a 2 (Valence: neutral vs. negative) by 2 (Item Type: object vs background) repeated-measures ANOVA. We found a significant main effect of Item Type, with higher recognition for objects than backgrounds, *F*(1, 51) = 179.80, *p* < .001. Critically, a significant Valence × Item interaction was observed, *F*(1, 51) = 30.75, *p* < .001. Specifically, negative scenes were associated with enhanced memory for objects but reduced memory for backgrounds relative to neutral scenes, consistent with an emotional tradeoff effect reported in previous studies^5,37^. There was no main effect of Valence (*F*(1, 51) = 0.78, *p* = .380), indicating that overall recognition accuracy did not differ between neutral and negative scenes.

Using the same approach as used for the questionnaire data, we examined whether neural affective patterns during sleep were related to individual differences in performance in the emotional tradeoff memory task across participants. Tradeoff was quantified by comparing the object-background differences in memory across valences (i.e., a higher tradeoff score would reflect that items were remembered better than backgrounds, for negative scenes more than for neutral ones; high scores therefore reflect a stronger bias toward negative information). We then correlated this behavioral measure with the wake-time-resolved affective activation profiles derived from cross-state decoding during sleep, using the same time-resolved, cluster-based permutation framework applied to the questionnaire analyses. An early wake-time decoding window, overlapping with the one observed for BDI-II and pre-sleep POMS, was also associated with memory revealed by the emotional tradeoff task. Specifically, stronger negative-valence decoding probabilities trained in this window (398–618 ms) were associated with a reduced overall tradeoff effect (*p_corrected_* = .039; *r* = -0.40, *p_uncorrected_* = .003; Fig 4b and Supp Fig 4d). This negative association suggests that individuals showing more negatively valenced patterns during sleep showed diminished emotional modulation of memory. Lower tradeoff scores may reflect a reduced sensitivity to negative relative to neutral stimuli or a generalized impairment in emotional differentiation.

Although tradeoff scores were not significantly correlated with BDI-II scores (*r* = -0.22, *p* = .120) or pre-sleep POMS scores (*r* = -0.19, *p* = .186), direct comparison of dependent correlations indicated that the association between tradeoff and negative-valence decoding probabilities (averaged over the overlapped time window across the three behavioral measures: 398–596 ms, *r* = -0.39) did not significantly differ from those involving BDI-II (*t*(49) = -1.30, *p* = .199) or pre-sleep POMS (*t*(49) = -1.44, *p* = .155). However, in a multiple regression analysis including BDI-II, pre-sleep POMS, and negative-valence decoding probabilities as simultaneous predictors, cross-decoding remained a significant predictor of tradeoff score (β = -0.36, *t*(48) = - 2.39, *p* = .021), whereas neither BDI-II (β = -0.04, *t*(48) = -0.24, *p* = .810) nor pre-sleep POMS (β = -0.02, *t*(48) = -0.14, *p* = .890) uniquely predicted tradeoff score. These findings indicate that the association between offline affective pattern expression and memory is not solely attributable to depressive symptom severity or mood state, consistent with a partially independent relationship.

Exploratory sleep-stage-specific analyses revealed that this negative association was significant during Stage 1 of sleep (*p_corrected_* = .013, during 376–618 ms; *r* = -0.44, *p_uncorrected_* < .001), Stage 2 of sleep (*p_corrected_* = .045, during 398–618 ms; *r* = -0.40, *p_uncorrected_* = .003), and pre-sleep or mid-sleep wakefulness (*p_corrected_* = .009, during 376–640 ms; *r* = -0.48, *p_uncorrected_* < .001; no significant clusters formed with α = 0.05 in other stages; Supp Fig 3f).

Finally, we examined whether the results obtained during the nap were sleep-specific. Participants completed two resting conditions (eyes-open and eyes-closed) in between the blocks of the affective localizer task. We repeated the cross-state classification analysis using the EEG data from these pre-sleep resting-state recordings (Supp Fig 5a). Results revealed an early BDI-II-linked cluster in both eyes-open (380–578 ms, *p_corrected_*= .015; *r* = 0.47, *p_uncorrected_* < .001) and eyes-closed resting-state (380–578 ms, *p_corrected_* = .043; *r* = 0.42, *p_uncorrected_* = .002; Supp Fig 5b). These clusters overlapped with the early cluster observed in the sleep data. A later significant cluster was identified during eyes-closed resting-state recordings (798–1062 ms, *p_corrected_*= .027; *r* = 0.43, *p_uncorrected_* = .002), but not during eyes-open recordings. This cluster overlapped with the second BDI-II-linked cluster identified during sleep. When examining the BDI subscales correlations, however, no significant clusters were found (*p*s*_corrected_* > .194 or no cluster formed with α = 0.05; Supp Fig 5c-d), suggesting either that subscale-specific dissociations emerge preferentially during sleep, or that the data collected during these short resting-state recordings did not yield sufficient statistical power to detect these correlations.

In contrast to the results obtained for BDI-II, we didn’t find any significant clusters when correlating the pre-sleep POMS with eyes-open nor eyes-closed resting-state data (*p*s*_corrected_* > .331, Supp Fig 5e). These results are in line with the finding that pre-sleep POMS was not significantly associated with affective pattern expression during pre-sleep or mid-sleep wakefulness. Together, these results suggest that pre-sleep mood may be preferentially associated with sleep relative to wakeful rest. Regarding emotional tradeoff, early clusters were observed in both eyes-open (380–600 ms, *p_corrected_* = .032; *r* = -0.42, *p_uncorrected_* = .002) and eyes-closed (380–622 ms, *p_corrected_* = .016; *r* = -0.48, *p_uncorrected_*< .001; Supp Fig 5f) resting-state recordings. The correlations followed the direction observed during sleep (i.e., more activation linked with less tradeoff). Both time windows closely overlapped with the decoding interval that predicted tradeoff during sleep, indicating that the relationship between affective pattern expression and emotional memory bias generalized across sleep and wakeful rest. Together, these findings suggest that neural affective patterns associated with stable, trait-like individual differences – including depressive symptom severity and interindividual differences in the preferential processing of emotional memories – are expressed across offline states, whereas associations with momentary mood state may be selectively expressed during sleep but not wakeful rest.

## Discussion

Affective disorders are tightly interlinked with sleep, but little is known about the mechanisms through which sleep supports emotional well-being. Here, we used a computational approach and neuroimaging data to investigate whether affective neural states during sleep are linked with trait- and state-level affective measures. Specifically, we combined time-resolved multivariate decoding with cross-state classification to investigate how affective traits and states are associated with spontaneous activation of valence-related neural patterns during sleep. Emotional valence was robustly decodable from EEG data during wakefulness. Critically, valence-related neural patterns re-emerged during sleep and wakeful rest, and the expression of negatively valenced neural patterns was associated with both state- and trait-level affective measures. Participants with more severe depression symptoms and more negative moods showed more negatively valenced neural patterns during offline states.

One explanation for the observed results is that information processing by affective networks during sleep is negatively biased. During wakefulness, negative biases in emotional processing (e.g., during ruminations) are a hallmark of affective disorders^26,38,39^. Excessive ruminations lead to prolonged depression episodes and predict future depressed mood^40,41^. Neuroimaging studies have similarly reported heightened default-mode and internally directed processing in depression, alongside increased rumination and negative self-referential thought^42–44^. Cognitive models of depression emphasize internally oriented negative biases that may remain latent during externally guided behavior but emerge during unconstrained mental states^42,43^. Our findings suggest that these biases in processing were perpetuated during sleep, potentially playing a role in the progression of MDD. While sleeping, previously encoded affective representations are spontaneously reactivated and reorganized through largely non-conscious neural processes. We suggest that negative biases in affective processing during sleep – just like during wake – impact waking mental health. For example, overnight non-conscious reactivation of mundane memories may be negatively biased, solidifying their negative interpretation. Notably, acute sleep deprivation can transiently alleviate depressive symptoms in patients with MDD^44^, suggesting that sleep-specific emotional processing may, in some cases, reinforce maladaptive affective patterns, in support of our proposed interpretation. However, as our current results are correlational, the causal role of affective processing during sleep remains to be explored.

The idea that emotional or neurocognitive processing during sleep may contribute to depressive symptoms has been proposed in prior theoretical and empirical work^45–47^. However, the direction of the observed correlation (i.e., demonstrating that negatively biased processing is linked with worse mental health status) seemingly goes against a long-held notion stating that sleep serves to dismantle memories for negative events. The “sleep to remember, sleep to forget” hypothesis states that during sleep (and particularly REM sleep) the negative affective tone of a memory is divorced from its declarative aspects, thereby rendering it less distressing. At face value, this hypothesis predicts that biases toward negative information processing during sleep would be adaptive and lead to better mental health outcomes. Of note, however, this process is thought to occur exclusively during REM sleep, whereas our findings are only significant for non-REM stages and wakefulness. In fact, our data revealed a distinct temporal cluster, observed only during REM, for which negatively valenced activation indeed correlated with fewer symptoms, in support of the “sleep to remember, sleep to forget”. Our data, collected over an afternoon nap, under-represented REM sleep, which is more prominent during nocturnal sleep, limiting statistical power (i.e., only 21 of the 52 participants entered REM at all). Future studies of nocturnal sleep physiology throughout the night could separately examine the relationship between symptoms and the affective activation patterns emerging during non-REM and REM sleep, shedding light on the role of both in offline emotional processing.

Decomposing depressive symptoms into cognitive-affective and somatic-performance dimensions further revealed a dissociation in how these symptom clusters mapped onto affective activity patterns during sleep. During wakefulness, we trained multiple neural classifiers which putatively reflect different neural representations of affective states. During sleep, distinct clusters linked with distinct representations were each associated with the different depression sub-scales: the cognitive-affective subscale of BDI-II, which measures emotional and mental components of depression^34,48^, and the somatic-performance subscale, which measures physical issues like fatigue or weight loss^34,48^. This temporal and state-dependent dissociation reflects heterogeneous alterations in affective dynamics that unfold across time and state^49^. Similarly, although most affective states were ubiquitously reflected across sleep stages and during wakeful rest, some state-specific difference emerged. However, it is not entirely clear whether these differences reflect differential statistical power or true categorical distinctions.

Several limitations warrant consideration. First, the present sample did not span all demographic parameters and consisted of young adults with predominantly low-to-moderate levels of depressive symptoms. The majority of participants (36 out of 52) scored in the minimal-to-mild range (BDI-II <= 13). While this allowed us to examine affective biases along a dimensional continuum in a non-clinical population, it remains an open question whether the observed patterns generalize to individuals with clinically diagnosed depression or more severe symptom profiles. Second, as mentioned above, the sleep opportunity consisted of a daytime nap rather than overnight sleep. As a result, relatively few participants entered REM sleep, limiting statistical power for REM-specific analyses and constraining inferences about REM-related affective processes. Moreover, affective dynamics observed during a nap may not fully capture the processes that unfold across a complete night of sleep, including interactions between successive sleep cycles. Future studies incorporating overnight recordings will be critical for determining whether the observed trait- and state-related affective patterns persist, evolve, or are transformed across extended sleep periods. Finally, the present findings are based on a single recording session per participant, precluding strong conclusions about the long-term stability of these affective patterns. Although associations with trait measures suggest a degree of stability, it remains unknown whether the observed sleep-based affective representations constitute a reliable individual “affective fingerprint”. Longitudinal and repeated-measures designs will be essential to determine whether these patterns are stable within individuals over time, sensitive to symptom change, or modifiable through intervention.

Taken together, our findings suggest that sleep and wakeful rest provide critical windows for revealing persistent affective biases that may remain latent during goal-directed wake activity. Standard methods for studying sleep’s role in mental health consider sleep’s basic structure (e.g., prevalence of different stages). Unfortunately, this approach is akin to judging the quality of an art museum based on the floor plan of the museum building – informative about structure, but silent about the content within. Here, we presented a novel approach using a unique cognitive-neuroscience-based framework to peer into the sleeping brain and understand the mechanisms through which sleep supports well-being, paving the way for further research into maladaptive neural processing in psychiatric disorders. Although our results are correlational, they fundamentally inform understanding of the role of sleep in affective processing, setting the stage for additional work examining and manipulating affective processing during sleep using machine-learning techniques and causal techniques in humans and non-human models.

## Methods

### Participants

Fifty-two participants (36 women, 15 men, 1 preferred not to state; mean age = 20.02, range: 18-25) from the University of California, Irvine campus community participated in the study for monetary compensation or class credits. All were right-handed and had normal or corrected-to-normal vision. Exclusion criteria included inability to understand English, history of neurological disease, history of major psychiatric disorder requiring hospitalization, history of major sleep disorders, and uncorrected hearing, visual or motor impairments. Participants were instructed to avoid alcohol and recreational drugs for 24 hours before the study, avoid caffeine consumption on the morning of the study, and wake up an hour earlier than their usual time. The experimental protocol was approved by the IRB of the University of California, Irvine and all participants provided written informed consent prior to the study.

### Material

All tasks were presented on a 24-inch capacitive touchscreen monitor (Dell P2418HT; Dell Inc., Texas, USA) with a native resolution of 1920 × 1080 pixels and a 60-Hz refresh rate. Participants were seated at a fixed viewing distance of approximately 60 cm. Experimental tasks were implemented in MATLAB R2022b (MathWorks Inc., Massachusetts, USA) using Psychtoolbox-3 for stimulus presentation and response collection. Responses were recorded via a standard keyboard.

Stimuli for the affective rating and localizer tasks consisted of 97 images selected from the International Affective Picture System (IAPS)^50^ and 5 images chosen from other online sources.

Images were chosen to form two valence categories (positive and negative) while equalizing arousal levels based on normative ratings (Fig 1b). Valence and arousal scores for the 48 negatively valenced IAPS images were 2.74 ± 0.44 (mean ± SD) and 4.83 ± 0.52 (mean ± SD), respectively. Valence and arousal scores for the 49 positively valenced IAPS images were 7.21 ± 0.44 (mean ± SD) and 4.82 ± 0.52 (mean ± SD), respectively. Each image was embedded within a uniform gray rectangular frame. The gray background served to equalize overall luminance and reduce variability in peripheral visual input across images. All images were resized to a uniform resolution of 960 x 540 pixels and were presented centrally on the screen.

Materials for the emotional tradeoff memory task were adapted from prior work examining emotion–memory interactions^37^. Stimuli consisted of composite scenes in which a single foreground object was superimposed onto a neutral background scene. The object in each trial was either neutral (e.g., a car parked on the street) or negatively valenced (e.g., a car accident on the street). A total of four counterbalanced stimulus sets were created for the encoding phase of the task. Each set included 40 composite images, comprising 20 scenes with a negative object and 20 scenes with a neutral object. Each participant was presented with one of the sets in a counterbalanced manner. For each background scene, two versions existed – one paired with a negative object and one paired with a neutral object – but each participant was exposed to only one version of each background during encoding. Participants were randomly assigned to one of the four stimulus sets, ensuring that object-background pairings were counterbalanced across participants.

For the recognition test following the nap, a total of 120 stimuli were presented, including objects and backgrounds tested separately. Object recognition included 60 stimuli (30 negative, 30 neutral), comprising 20 previously seen objects (“old”), 20 perceptually similar lures, and 20 novel objects. Background recognition likewise included 60 stimuli, all neutral in valence: 20 old backgrounds (previously paired with either negative or neutral objects), 20 similar backgrounds, and 20 novel backgrounds. This design allowed assessment of memory for objects and backgrounds independently, as well as quantification of emotional tradeoff effects reflecting differential prioritization of emotional versus contextual information.

### Experimental Procedure

Participants arrived at the laboratory around 1 pm and were fitted with a 64-channel EEG cap. They then completed an affective rating task in which they evaluated 102 positive and negative images on a 1-9 Likert scale, with higher values indicating more positive valence. The images were presented in pseudorandom order, so that no more than four images of the same valence class would be presented in sequence. Participants were also presented with negative and positive videos, which were not included in the analysis and are therefore not described in more detail. After rating all images, the same stimuli were used in an affective localizer task conducted with continuous EEG recording. In the affective localizer task, five blocks of all 102 images were presented, with all images presented in each block in a pseudorandom order so that no more than four images of the same valence class would be presented in sequence. On each trial, participants viewed an image and pressed the down arrow key on the keyboard when they were ready to report its valence (self-paced response with up to 10 s to proceed). The image was then replaced by either a ‘Happy?’ or ‘Sad?’ prompt (randomized across repetitions and balanced across valance class), and participants responded using left/right arrow keys (‘no’/‘yes’, respectively; self-paced response with up to 3 s to proceed). For example, if a negatively valenced image was followed by a ‘Happy?’ prompt, a correct response would have been ‘no’, indicated by a left-key choice. Note that since prompts were balanced across valence classes, this design does not confound motor action with valence (i.e., neural classification does not reflect motor execution). Of the 102 images for each block, the trials involving the first two images (a positively valenced one and a negatively valenced one) were not included in the analysis.

Between the first and second blocks of the task, participants were asked to take a 3-minute break, during which they were asked to stare at an on-screen crosshair with their eyes open. Between the second and third blocks, participants were asked to take a 3-minute break, during which they were asked to close their eyes. A single beep sound instructed them to open their eyes and continue to the next block. After completing all blocks, participants completed the first phase of the emotional tradeoff memory task^5^. This task included an initial incidental encoding portion, followed by a test after the nap. During the training phase, participants viewed images depicting an object superimposed on a neutral background. Participants rated how likely they would be to approach each scene on a 1–9 Likert scale, encouraging affective engagement with the stimuli without explicitly anticipating the subsequent memory test.

Prior to the nap, participants completed POMS^27^, STAI-S^28^, and SSS^29^. They were then provided with a 90-minute opportunity to sleep on a comfortable bed in the laboratory while EEG was continuously recorded. Upon awakening, SSS, POMS and STAI-S were re-administered, followed by a surprise memory test in which recognition memory of the emotional tradeoff task was assessed separately for objects and backgrounds. In total, 120 images were presented and participants had to indicate for each whether it was previously presented (‘old’), new, or similar but not identical to a previously presented image (‘similar’). Sixty of the images were of objects, including thirty negatively valenced and thirty neural images. The sixty images included twenty old images, twenty similar images, and twenty neutral images. The remainder sixty images were of backgrounds. All backgrounds were neutral. Twenty were novel and not presented during training, twenty were presented beforehand (half with negatively valenced and half with neutral objects), and twenty were similar to a background presented during training (half with negatively valenced and half with neutral objects). Trials were presented in a random order and all responses were self-paced.

Finally, participants completed a set of trait questionnaires, including BDI-II^32^, PHQ^33^, the Anxiety Sensitivity Index (ASI)^51^, the Mood and Anxiety Symptom Questionnaire (MASQ)^52^, and the Questionnaire of Unpredictability in Childhood (QUIC)^53^. Item 9 (“Suicidal thoughts or wishes”) of BDI-II was excluded in the current study, as no protocol for dealing with suicidality was implemented to deal with potential adverse consequences. All BDI-II related analyses were based on the remaining 20 items. After completing all tasks, participants removed the EEG cap and electrodes, cleaned up, and were then debriefed, compensated, and dismissed.

### EEG Data Acquisition and Preprocessing

EEG was recorded from 64 scalp electrodes positioned according to the 10-20 system using the eego system (ANT Neuro, Netherlands), with a sampling rate of 500 Hz. Impedances were kept below 20 kΩ. EEG data was preprocessed using the Fieldtrip toolbox in Matlab (R2023a). The continuous EEG data of the affective localizer task was filtered between 0.5-35 Hz using a band-pass Butterworth filter, and re-referenced to the average of the two mastoid electrode channels. The EMG channel was excluded for wake EEG data, and bad electrodes were interpolated. Independent Component Analysis was conducted after down-sampling to 250 Hz on all non-interpolated EEG channels (excluding reference, EMG, and EOG). Independent component reflecting artifacts (ocular, muscle) were identified through topographies and component time courses and removed. Manual artifact rejection was performed to exclude high-amplitude noise and transient artifacts. The resulting cleaned continuous data were then segmented into epochs time-locked to stimulus onset from -2 to 3 s.

Sleep EEG during the 90-min nap was preprocessed similarly, except for a few modifications appropriate for sleep physiology: the EMG channel was preserved and filtered at 10-100 Hz. Independent Component Analysis was not performed on sleep EEG. Cleaned continuous sleep data were used for sleep staging and cross-state decoding analysis.

### Sleep Staging

Sleep architecture during the 90-min nap was scored by two independent raters following the American Academy of Sleep Medicine criteria^54^. Staging was performed on 30-s epochs of the preprocessed continuous EEG, EOG, and EMG signals. Each rater assigned stages (N1, N2, N3, REM, Wake) based on standard polysomnographic features. Inter-rater agreement was assessed across all epochs, and disagreements were resolved. Sleep measures and stages were derived from the consensus scoring and used in subsequent analyses. We did not exclude any participants based on their sleep architecture.

### Decoding Analysis During Wake

We trained time-resolved within-participant multivariate classifiers on EEG data obtained from the affective localizer task to distinguish positively valenced and negatively valenced images. For each participant, we used a 5-block cross-validation approach repeated over 20 iterations. Data were first averaged within sliding time windows of 102 ms moved in steps of 22 ms across the trial time-course. We used linear support vector machines (SVMs) implemented in MATLAB’s *fitcecoc* function. This procedure yielded a temporal decoding accuracy curve for each participant.

Statistical significance of wake decoding accuracy was assessed using a cluster-based permutation test to control for multiple comparisons across time. For each of 1,000 permutation, class labels were randomly assigned at the trial level within each participant, block and iteration, preserving the temporal structure. Decoding accuracy was recomputed at each time point, averaged across iterations and blocks, and smoothed using a five-point moving average, yielding a time-resolved accuracy trace for each participant. Then, decoding accuracy was compared against chance on the group level at each time point using a one-sample *t*-test. Temporally contiguous time points exceeding the cluster-forming threshold (two-sided α = 0.05) were grouped into clusters, and a cluster-level statistic was computed as the sum of *t*-values within each cluster. For each random permutation, the maximum cluster-level statistic was retained to form a null distribution. Observed clusters were considered significant if their cluster statistics exceeded the 95^th^ percentile of the permutation null distribution. Cluster-corrected *p* values were computed as the proportion of permutation-derived clusters that matched or exceeded the observed cluster.

### Cross-state Decoding During Nap and Resting-State Recordings

Classifiers trained during wakefulness were applied to the sleep data. EOG and EMG electrodes were dropped for this analysis. Sleep EEG data were sampled every 10 seconds throughout the 90-minute nap. For each time point, the data were averaged over a 250-ms time span. These sleep samples constituted the test data for cross-state decoding. The classifiers trained timepoint-by-timepoint on the wake EEG datasets were applied to the nap data. Specifically, for each time point relative to image onset during wakefulness, we used all trained SVMs (i.e., one per time point) to predict class value (i.e., 1 for positive and 2 for negative) for each sleep sample (Fig 3a). The prediction output values (1 or 2) were averaged across the models created for the 5-fold cross validation blocks and 20 iterations (100 models total per wake time point per participant). The result was a time-resolved measure of the frequency of negative pattern activation during sleep, with a single data point for the intersection of each wake time point and sleep time point. The values ranged between 1 to 2, with a larger value meaning higher probability of negative activation at a specific sleep sample. This produced a 2-D matrix of cross-state decoding per participant (training-time × sleep-time, Fig 3c, top panel).

A similar approach was used to calculate affective neural activation during resting-state recordings, calculated separately for periods of time with eyes open and eyes closed. Resting-state EEG was collected during the affective localizer session and preprocessed together with the localizer EEG. The data was sampled every 2 seconds to obtain ∼75 sampled events. Similar to the sleep decoding, for each sampled event, the data was averaged over a 250 ms time span. These event-level representations constituted the test data for cross-state decoding during resting-state recordings, followed by the same classification and averaging procedures as in the sleep analysis. Due to technical issues, EEG data from two participants were missing from the eyes-open resting-state recordings (*n* = 50), from one from the eye-closed resting-state recordings (*n* = 51).

To quantify the negative activation probability across the entire nap or resting-state period, as well as within specific sleep stages, designated classes for all test timepoints were averaged to obtain the mean affective activation frequency (a value ranging from 1 to 2, with larger value meaning higher probability of negative activation) at each specific wake-training time point. For example, this strategy was used to collapse values across all time points during sleep or to collapse values across all time points included in a certain sleep stage. Data were only collapsed over sleep or resting-state time, while the wake localizer trial-based temporal dynamics were preserved.

### Time-resolved Correlation Analysis of Questionnaires Measures

Our analytic approach was to link affective activation patterns to state- and trait-level wakeful affective measures. We assessed the relationship between affective activation during rest and behavioral measures using a cluster-based permutation correlation test of 1,000 permutations. First, for each training time point (locked to stimulus onset during wakefulness), we computed Pearson’s *r* between behavioral measures and the affective activation values across participants. Time points with *p* < .05 (uncorrected) were grouped into contiguous clusters. Next, 1,000 permutations were performed by randomly shuffling the behavioral measures across participants (the reassigned level was the same across time points for each participant in each permutation, preserving participant-level temporal dynamics). For each permutation, the correlation and cluster statistics were recomputed, creating a null distribution of the largest contiguous clusters across participants for each permuted dataset. For each real cluster, we calculated a corrected *p*-value based on the summed *t*-value for the real clusters and the distribution of max summed *t*-values across permutations (two-tailed analysis). A cluster was considered significant if its corrected *p* < .05.

Using this analytical framework, we examined whether individual differences in affective and behavioral measures were related to neural affective activation patterns derived from cross-state decoding. Behavioral measures included depressive symptom severity assessed with the BDI-II questionnaire. Consistent with the widely used two-factor structure^34^, we computed two subscale scores of BDI-II: Cognitive-Affective subscale (items 1-8, 10-11, and 14) and Somatic-Performance subscale (items 12, 13 and 15-21). Each subscale score was obtained by summing the corresponding item scores, and the total BDI-II score was computed as the sum of all 20 items. Higher scores indicate greater depressive severity. In addition to the BDI-II scores, the pre-sleep and post-sleep mood states were assessed with the POMS questionnaire.

For each measure, we correlated individual difference scores with wake time-resolved affective decoding profiles using a time-resolved, cluster-based permutation approach. Neural affective measures were derived from cross-state decoding outputs and included decoding values averaged across the entire nap period, decoding values averaged within specific sleep stages (N1, N2, N3, and REM), and decoding values obtained from resting-state recordings, analyzed separately for eyes-open and eyes-closed conditions. For analyses involving averaged neural measures (e.g., across the entire nap, within a sleep stage, or across resting-state epochs), values were collapsed only across the resting period/stage, while preserving the wake training time dimension of the decoding results. This procedure yielded wake time–resolved correlation time courses for each behavioral measure and neural condition, which were evaluated for statistical significance using the cluster-based permutation testing explained above to control for multiple comparisons across time.

### Analysis of the emotional tradeoff memory task

Emotional tradeoff memory was assessed using an established object-background paradigm in which recognition memory for neutral background elements is modulated by the emotional valence of concurrently presented objects. For previously studies (“old”) items, correct recognition was defined as endorsement of either “old” or “similar” responses. For each participant, an overall tradeoff score was computed as: (recognition accuracy for negative objects minus recognition accuracy for neutral objects) minus (recognition accuracy for backgrounds paired with negative objects minus recognition accuracy for backgrounds paired with neutral objects). Positive values therefore indicate enhanced memory for negative objects combined with reduced memory for their associated backgrounds, consistent with an emotional tradeoff effect. To test whether individual differences in emotional tradeoff were related to affective neural activation during sleep, we correlated each participant’s overall tradeoff score with their cross-state decoding time series (obtained as described above). Specifically, Pearson’s *r*s were calculated between the tradeoff scores and the decoding time series across participants at each wake training time point. Statistical significance was assessed using a cluster-based permutation testing (α = 0.05) like before, with clusters defined as contiguous time points exceeding an uncorrected threshold and evaluated against a permutation-derived null distribution, controlling for multiple comparisons across time while preserving sensitivity to temporally extended effects.

## Acknowledgements

This project was funded by the National Institute of Health (award # P50 MH096889 & DP1 HL179370). The authors would like to thank Neda Morakabati, Taryn Mallory, Charli Taylor. Evelyn Maivy Le, Maya Pourezza, and Pavana Upadhyaya for assistance with sleep scoring and initial data analysis. They also thank Elizabeth Kensinger for providing access to her lab’s stimulus database.

