## Supplementary Materials for "Depression symptoms are associated with affective neural processing during sleep and rest"

**Supplementary Figures**

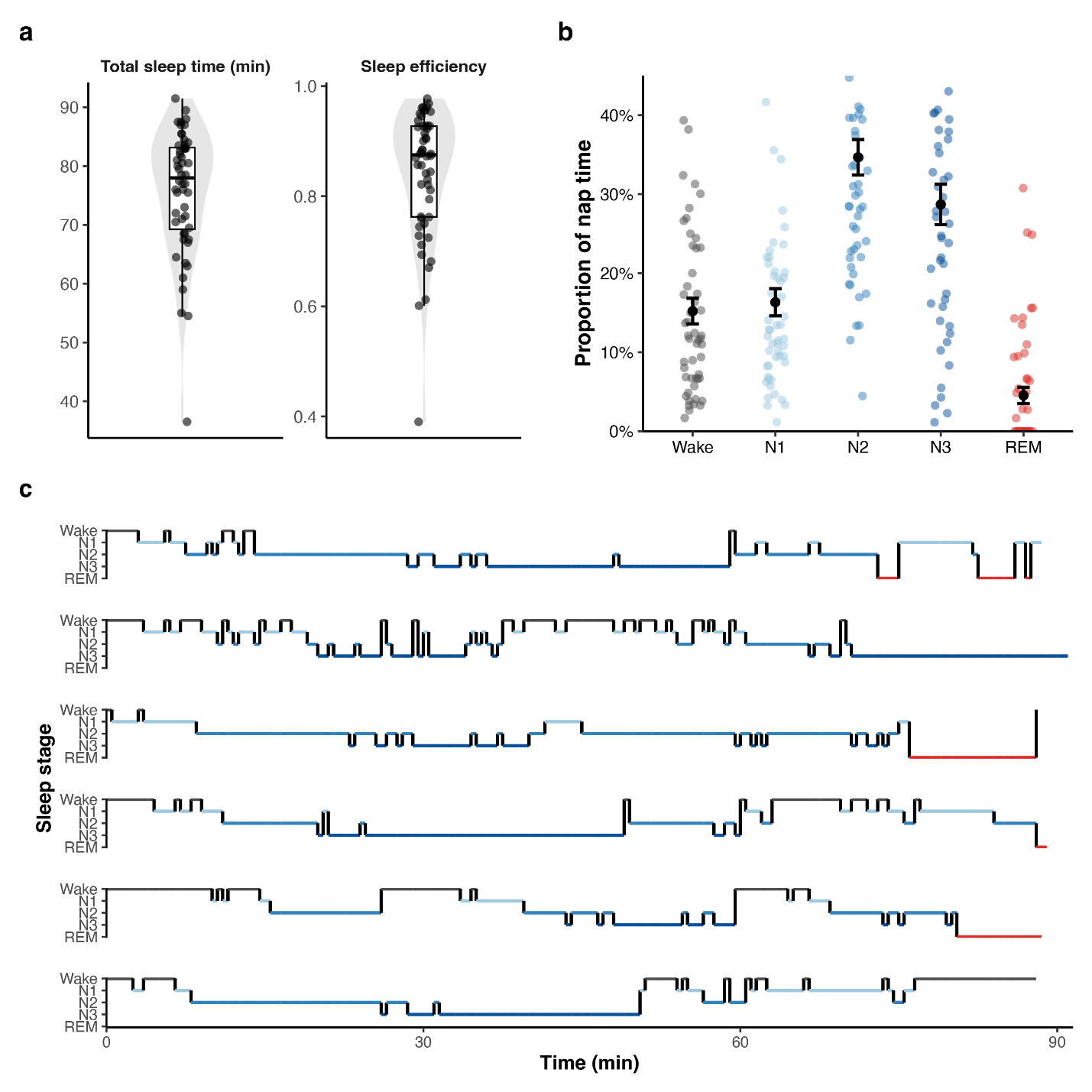

*Supplementary Figure 1. Sleep architecture*

*(a) Distribution of total sleep time and sleep efficiency (total sleep time/recorded sleep opportunity) across participants. Gray violin depicts the full distribution; dots represent individual participants; black boxplots show the median and interquartile range.*

*(b) Proportion of time spent in each sleep stage during the nap opportunity. Dots represent individual participants; black markers indicate mean ± s.e.m.
(c) Representative individual hypnograms illustrating typical nap sleep architecture. REM sleep epochs are highlighted in red. REM – rapid-eye-movement sleep; N1/N2/N3 – stages 1/2/3, respectively.*

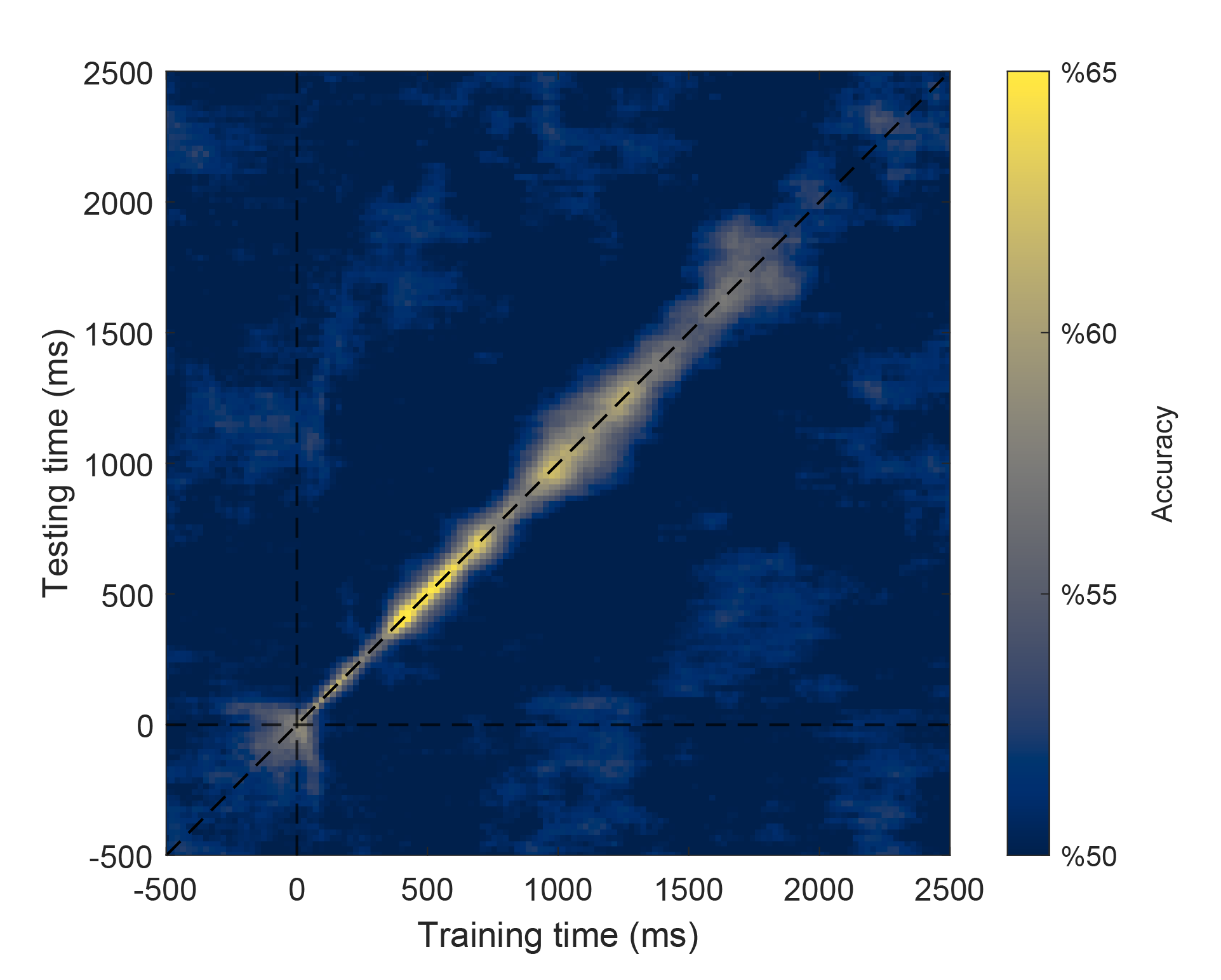

*Supplementary Figure 2. Wake-state decoding temporal generalization*

*Wake-state time generalization matrix showing classifier performance across training and testing time points locked to stimulus onset during trials of the affective localizer task. Data across time points during wakefulness were submitted to all models trained on all time points. The diagonal reflects the effects observed in Fig 2b. This analysis demonstrates that models trained on a specific time point do not generalize to other time points (i.e., each model leverages different neural patterns distributed across the scalp to accurately classify valence).*

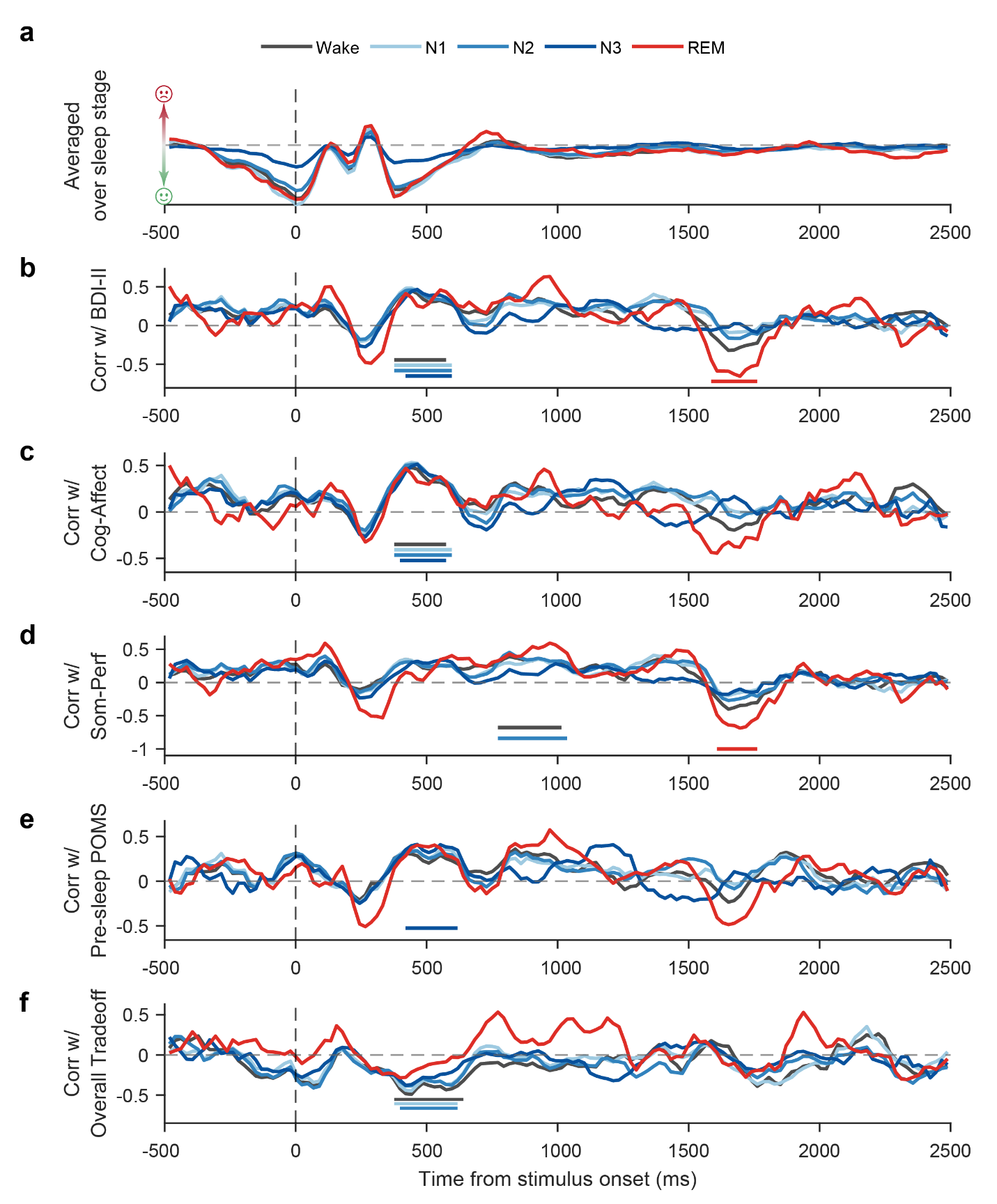

*Supplementary Figure 3. Sleep stage specific decoding*

*(a) Cross-state decoding time courses shown separately for each sleep stage (Wake, N1, N2, N3, REM). Values reflect average cross-decoding prediction, with positive values indicating higher probability of negative valence, and negative values indicating higher probability of positive valence.*

*(b) Stage-specific correlations between decoding strength and total depressive symptom severity (BDI-II total score) across participants.*

*(c) Stage-specific correlations between decoding strength and the BDI-II Cognitive–Affective (Cog-Affect) subscale.*

*(d) Stage-specific correlations between decoding strength and the BDI-II Somatic–Performance (Som-Perf) subscale.*

*(e) Stage-specific correlations between decoding strength and pre-sleep mood state (measured using POMS).*

*(f) Stage-specific correlations between decoding strength and the emotional tradeoff memory score [overall tradeoff, i.e., (negative object memory minus negative background memory) minus (neutral object memory minus neutral background memory)].*

*In all panels, time periods with significant correlations (p < 0.05, corrected) are marked by colored horizontal lines for each sleep stage. BDI-II – Beck Depression Inventory-II; POMS – Profile of Mood State questionnaire; REM – rapid-eye-movement sleep; N1/N2/N3 – stages 1/2/3, respectively.*

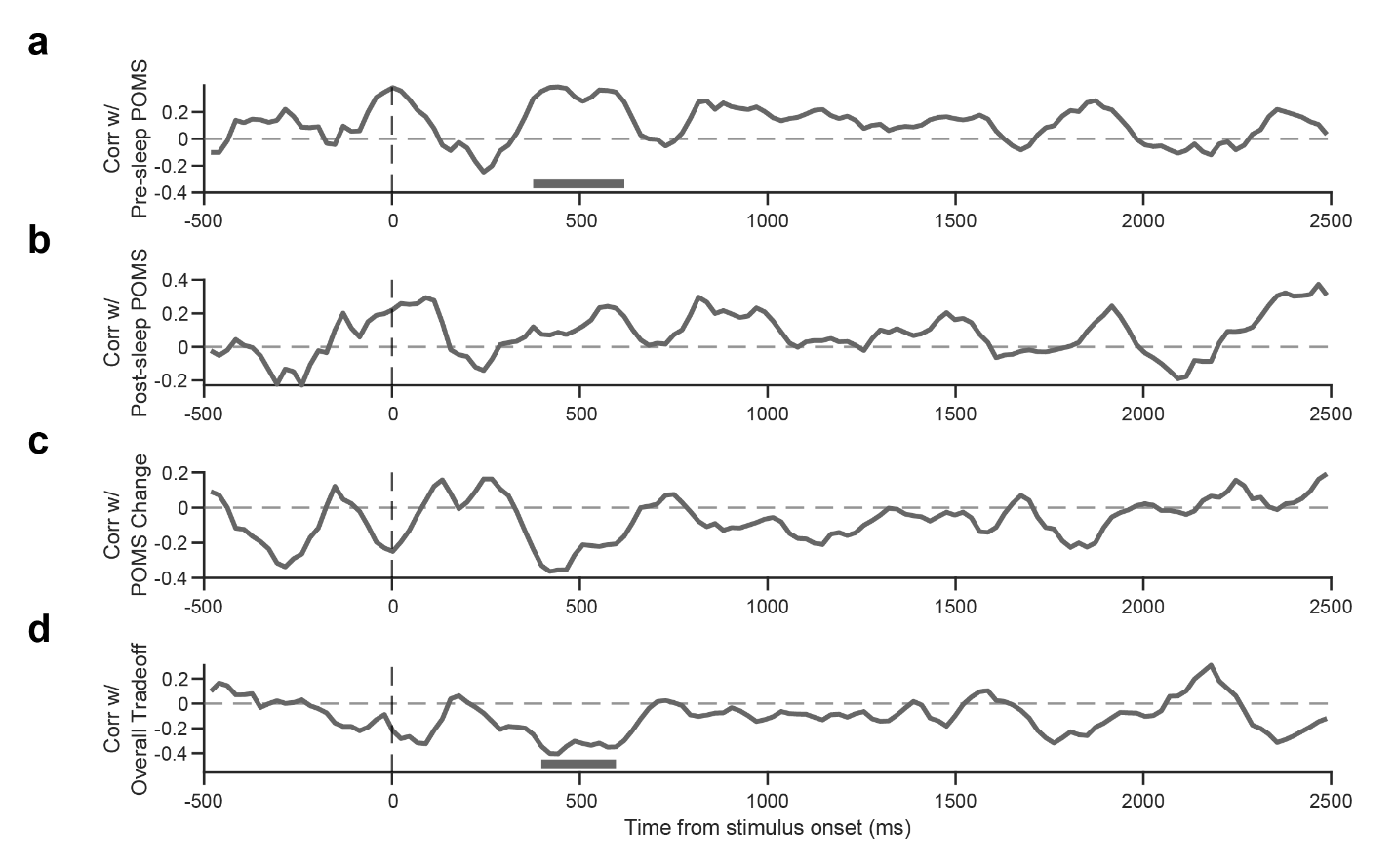

*Supplementary Figure 4.* *Cross decoding correlation time course with other behavioral measures*

*(a) Time-resolved correlations between cross-state decoding for the entire nap and pre-sleep mood state (measured using POMS).*

*(b) Time-resolved correlations between cross-state decoding for the entire nap and post-sleep mood state (measured using POMS).*

*(c) Time-resolved correlations between cross-state decoding for the entire nap and change in mood across the nap (post-sleep minus pre-sleep POMS).*

*(d) Time-resolved correlations between cross-state decoding for the entire nap and overall emotional memory tradeoff [overall tradeoff, i.e., (negative object memory minus negative background memory) minus (neutral object memory minus neutral background memory)].*

*In all panels, time periods with significant correlations (p < 0.05, corrected) are marked by horizontal lines. POMS – Profile of Mood State questionnaire.*

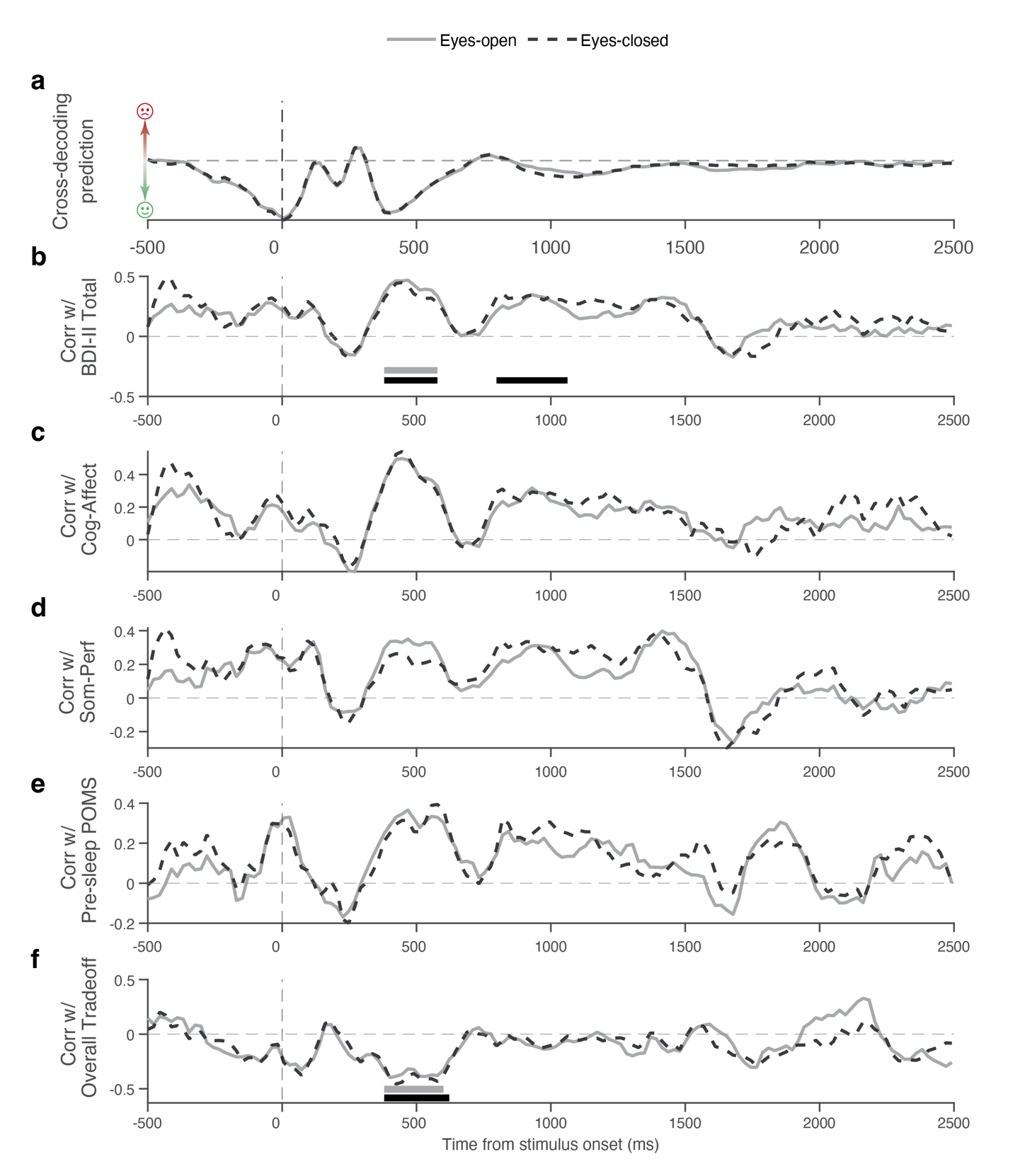

*Supplementary Figure 5. Resting state cross decoding correlation time course with behavioral measures*

*(a) Cross-state decoding time courses shown separately for eyes-open resting state and eyes-closed resting state. Values reflect average cross-decoding prediction, with positive values indicating higher probability of negative valence, and negative values indicating higher probability of positive valence.*

*(b) Time-resolved correlations between cross-state decoding for the resting states and total depressive symptom severity (BDI-II total score) across participants.*

*(c) Time-resolved correlations between cross-state decoding for the resting states and the BDI-II Cognitive–Affective (Cog-Affect) subscale.*

*(d) Time-resolved correlations between cross-state decoding for the resting states and the BDI-II Somatic–Performance (Som-Perf) subscale.*

*(e) Time-resolved correlations between cross-state decoding for the resting states and pre-sleep mood state (POMS).*

*(f) Time-resolved correlations between cross-state decoding for the resting states and overall emotional memory tradeoff [overall tradeoff, i.e., (negative object memory minus negative background memory) minus (neutral object memory minus neutral background memory)].*

*In all panels, time periods with significant correlations (p < 0.05, corrected) are marked by horizontal lines. BDI-II – Beck Depression Inventory-II; POMS – Profile of Mood State questionnaire.*

**Supplementary Tables**

Supplementary Table 1. Sleep architecture summary

Sleep stages were scored according to American Academy of Sleep Medicine (AASM) criteria. Values reflect the 90-min nap opportunity. Sleep-stage metrics are reported across all participants (*N* = 52); all participants entered N1 and N2 sleep, whereas 50 entered N3 sleep and 21 entered REM sleep.

| **Metric** | **Mean** ± **SD** | **Range** |
| --- | --- | --- |
| Total sleep time (min) | 75.69 ± 10.65 | 36.50–91.50 |
| Sleep efficiency (%) | 84.24 ± 11.67 | 39.04–97.75 |
| Wake (min) | 13.71 ± 10.69 | 1.50–56.50 |
| N1 (min) | 14.72 ± 11.21 | 1.00–58.00 |
| N2 (min) | 31.23 ± 14.56 | 3.50–74.00 |
| N3 (min) | 25.62 ± 16.27 | 0.00–70.50 |
| REM (min) | 4.12 ± 6.79 | 0.00–28.00 |

Supplementary Table 2. Questionnaire descriptive statistics

| **Questionnaire** | **Mean** ± **SD** | **Range** |
| --- | --- | --- |
| Anxiety Sensitivity Index (ASI) | 25.37 ± 13.39 | 3.0–60.0 |
| Beck Depression Inventory-II (BDI-II) | 10.83 ± 7.30 | 0.0–34.0 |
| Mood and Anxiety Symptom Questionnaire (MASQ) | 44.10 ± 8.92 | 27.0–76.0 |
| Patient Health Questionnaire (PHQ-9) | 5.73 ± 4.55 | 0.0–18.0 |
| Questionnaire of Unpredictability in Childhood (QUIC) | 9.19 ± 7.21 | 0.0–30.0 |
| State-Trait Anxiety Inventory, State questions only (STAI-S, pre-sleep) | 33.87 ± 7.53 | 23.0–55.0 |
| Profile of Mood State (POMS, pre-sleep) | 8.83 ± 9.24 | -13.0–32.0 |
| Stanford Sleepiness Scale (SSS, pre-sleep) | 4.10 ± 1.30 | 1.0–7.0 |
| State-Trait Anxiety Inventory, State questions only (STAI-S, post-sleep) | 30.60 ± 5.79 | 20.0–41.0 |
| Profile of Mood State (POMS, post-sleep) | 0.42 ± 6.35 | -14.0–17.0 |
| Stanford Sleepiness Scale (SSS, post-sleep) | 2.60 ± 0.77 | 1.0–5.0 |

Supplementary Table 3. Correlations between depressive symptom severity and behavioral measures. Pearson correlations between BDI-II total score and all other subject-level behavioral measures in the study (two-tailed). Confidence intervals reflect 95% intervals for the correlation coefficient. *p*-values were corrected for multiple comparisons across the 14 tests using the Benjamini-Hochberg false discovery rate (FDR) procedure.

| **Measure** | ***r*** | **CI_low** | **CI_high** | ***p*** | ***p* (FDR)** |
| --- | --- | --- | --- | --- | --- |
| BDI Cognitive–Affective | 0.91 | 0.84 | 0.95 | < .001 | < .001 |
| BDI Somatic–Performance | 0.89 | 0.82 | 0.94 | < .001 | < .001 |
| PHQ-9 | 0.89 | 0.82 | 0.94 | < .001 | < .001 |
| ASI | 0.21 | -0.07 | 0.45 | .142 | .166 |
| MASQ | 0.37 | 0.11 | 0.58 | .007 | .014 |
| Pre-sleep POMS | 0.58 | 0.36 | 0.74 | < .001 | < .001 |
| Post-sleep POMS | 0.22 | -0.05 | 0.47 | .113 | .153 |
| POMS change (post–pre) | -0.46 | -0.65 | -0.22 | .001 | .001 |
| Pre-sleep STAI | 0.46 | 0.22 | 0.65 | .001 | .001 |
| Post-sleep STAI | 0.29 | 0.02 | 0.52 | .039 | .064 |
| STAI change (post–pre) | -0.28 | -0.51 | -0.01 | .041 | .064 |
| Emotional trade-off | -0.22 | -0.46 | 0.06 | .120 | .153 |
| Negative valence rating | 0.19 | -0.09 | 0.44 | .186 | .200 |
| Positive valence rating | -0.09 | -0.35 | 0.19 | .535 | .535 |
